# Erythroid differentiation displays a peak of energy consumption concomitant with glycolytic metabolism rearrangements

**DOI:** 10.1101/514752

**Authors:** Angélique Richard, Elodie Vallin, Caroline Romestaing, Damien Roussel, Olivier Gandrillon, Sandrine Gonin-Giraud

**Affiliations:** Univ Lyon/ENS de Lyon/Univ Claude Bernard/CNRS UMR 5239/INSERM U1210, Laboratory of Biology and Modelling of the Cell, Lyon, France; UMR 5023 CNRS/Université de Lyon, Laboratoire d’Ecologie des Hydrosystèmes Naturels et Anthropisés, Lyon, France; Inria Team Dracula, Inria Center Grenoble Rhône-Alpes, Grenoble, France

## Abstract

It has been suggested that a switch from glycolysis, with lactate production, toward mitochondrial oxidative phosphorylation (OXPHOS) could be a driving force during stem cell differentiation.

Based upon initial results, from our previous work, showing a drop in LDHA mRNA level during the differentiation of chicken erythroid progenitors, we studied metabolism behavior to question whether such switch might also be operating in those cells.

We first analyzed the level of 9 enzymes, including LDHA, involved either in glycolysis or OXPHOS, in self-renewing and differentiating cells. Our results suggest that erythroid differentiation might be accompanied by an enhancement of the respiratory chains and glycolysis activities at 12h, followed by a strong decline of the glycolytic pathway and a stabilization of OXPHOS.

To confirm that OXPHOS might be increased and glycolysis decreased during erythroid differentiation, we measured lactate concentration and mitochondrial membrane potential (MMP) of self-renewing and differentiating cells. Our findings show that at 12h-24h of differentiation, a surge of energy is needed, which could be fueled jointly by glycolysis and OXPHOS. Then the energy demand comes back to normal and might be supplied by OXPHOS instead of lactate production through glycolysis.

These results support the hypothesis that erythroid differentiation is associated with a metabolic switch from glycolysis to OXPHOS.

We also assessed LDHA role in erythroid progenitors self-renewal and the metabolic status changes. Inhibition experiments showed that LDHA activity could be involved in the maintenance of erythroid progenitors self-renewal, and its decline could influence their metabolic status.

Finally, we investigated whether these metabolic rearrangements were necessary for erythroid differentiation. The addition of an inhibitor of the respiratory chains affected progenitors ability to differentiate, suggesting that the metabolic switch from glycolysis toward OXPHOS might act as a driving force for erythroid differentiation.

**Author summary:** Single-cell based gene expression data from one of our previous publication pointed out significant variations of LDHA level, an important metabolism player, during erythroid differentiation. Deeper investigations highlighted that a metabolic switch occurred along differentiation of erythroid cells as previously emphasized in stem cell differentiation. More precisely our finding showed that self-renewing progenitor cells relied mostly upon a glycolytic, lactate-productive, metabolism and required LDHA activity, whereas differentiating cells, mainly involved the aerobic mitochondrial oxidative phosphorylation (OXPHOS). However our careful kinetic study demonstrated that these metabolic rearrangements were coming along with a particular temporary event, occurring within the first 24h of erythroid differentiation. The activity of glycolytic metabolism and OXPHOS rose jointly with ATP production at 12-24h of the differentiation process before lactate-productive glycolysis sharply fall down and energy needs decline. Finally, our results showed that the metabolic switch mediated through LDHA drop and OXPHOS upkeep might be necessary for erythroid differentiation. We also discuss the possibility that metabolism, gene expression and epigenetics could act together in a circular manner as a driving force for differentiation.

## Introduction

Metabolism is a biological process mostly involved in energy consumption, production and distribution, which is essential for cell functions and survival.

An emerging theme is the possible causal involvement of metabolic changes during a differentiation process. In particular, glycolysis was shown to be characteristic of self-renewing stem cells and of different types of progenitors, such as neuronal and hematopoietic progenitor cells [1, 2]. It has been demonstrated that stem cells tend to switch from glycolytic metabolism toward mitochondrial oxidative phosphorylation (OXPHOS) while they differentiate [3–5]. Accordingly, it was suggested that a balance between glycolysis and OXPHOS metabolism could somehow guide the choice between self-renewal and differentiation in stem cells fate [6]. It has also been demonstrated that preventing the shutting down of the glycolytic pathway was detrimental for neuronal differentiation [4]. Otherwise, regarding somatic cells reprogramming into induced pluripotent stem cells, it has been reported that pluripotency induction required the concomitant upregulation of glycolytic metabolism and downregulation of OXPHOS [7].

Regarding erythroid differentiation, progenitors depend on glycolysis and present a Warburg-like profile as they display a high proliferation rate and produce an abundant amount of lactate [8]. However, few is known about glucose metabolism behavior during erythroid differentiation, when compared to erythroid progenitors self-renewal. Mature mammals erythrocytes were shown to rely upon lactate dehydrogenase to fulfill their functions, depending therefore upon the glycolytic pathway [9, 10]. This is in agreement with the fact that mammalian erythroid cells totally eliminate their mitochondria at the end of their maturation [11]. Therefore, it is not possible for mammalian erythrocytes to produce ATP through mitochondrial OXPHOS anymore. However it is different for avian erythroid cells, that keep their mitochondria until the very last mature state [12].

Recent results from high-throughput transcriptome analysis highlighted that key actors of the glycolytic metabolism, such as LDHA, decreased while erythroid progenitors differentiate, both in chickens [13], and mice [14]. Moreover another recent study performed in humans at the proteome level emphasized similar behaviour of enzymes involved in the glycolytic pathway, during erythroid differentiation [15]. These new findings suggested that erythroid progenitors metabolism could exit from its glycolytic state during the differentiation process, and might switch toward OXPHOS.

In light of our previous work [13] and of these new insights we explored the hypothesis that erythroid progenitor cells could undergo a metabolic switch from glycolysis toward OXPHOS while they differentiate, and that this shift might be necessary for the differentiation process.

We therefore assessed different parameters of the glycolytic metabolism during the first three days of primary avian erythroid progenitor cells differentiation. We first analyzed the protein level of a few key enzymes involved either in glycolysis or OXPHOS pathway, at six time-points of the differentiation process. To confirm our results at the physiological level we then compared lactate production, indicating glycolytic rate, and mitochondrial membrane potential (MMP), that partially reflects OXPHOS activity, of self-renewing and differentiating cells. We further dissected OXPHOS mechanisms by analyzing cell respiration response to different respiratory chains inhibitors. Finally, LDHA role in erythroid progenitors self-renewal was assayed using known molecular inhibitors and the CRISPR/Cas9 knock-out technology. Then, to investigate whether LDHA downregulation could be responsible for metabolic state changes in erythroid progenitors, we measured MMP of self-renewing cells following inhibition of LDHA activity. Finally, we assessed whether or not the metabolic shift toward OXPHOS might be necessary for T2EC differentiation.

As a conclusion, our findings support our hypotheses suggesting that (1) erythroid maturation could be accompanied with a switch from glycolysis, resulting in lactate production, toward OXPHOS, (2) LDHA is involved in self-renewal maintenance, and (3) the metabolic shift toward OXPHOS might play an important role in erythroid differentiation.

## Materials and methods

### Cell culture conditions and treatments

T2EC were grown as previously described [16]. For LDHA inhibitors treatment, cells were cultured at 1.25.10^5^ cells/ml in fresh culture media containing either Galloflavin (50 *μ*M) (SIGMA), FX11 (30 *μ*M) (SIGMA) or DMSO (SIGMA). Thereafter, cells were counted every 24h, or harvested for lactate and ISX analysis.

For antimycin A treatment, cells were cultured at 1.25.10^5^ cells/ml in fresh differentiation media containing antimycin A (10 nM) (SIGMA) or DMSO (SIGMA). After 48h, cells were counted using trypan blue (SIGMA) or stained with Benzidine (SIGMA). For RTqPCR analysis following antimycin A or DMSO treatments, living cells were first sorted using a Lymphocyte Separation Medium (LSM) (SIGMA) and then harvested to extract RNA (for more details see subsection *Quantitative Real-Time Polymerase Chain Reaction assays*).

### Cell growth and differentiation measurements

Cell growth was evaluated by counting living cells using Trypan blue staining. Cell differentiation was assessed using benzidine staining (SIGMA) [17] and betaglobin mRNA level using RTqPCR (see subsection *Quantitative Real-Time Polymerase Chain Reaction assays* for descriptions).

### Cell transfection and sorting

Cells were transfected by electroporation using the Neon transfection system 100*μ*l kit (Thermo fisher) according to manufacturers recommendations. For each condition 10.10^6^ cells were washed once with 1X PBS. Cell pellets were resuspended in T transfection buffer supplied in the kit and 10*μ*g of pCRISPR-P2A-tRNA (see subsection *CRISPR plasmid constructions* for descriptions) was added. Cells were loaded into dedicated tips using the Neon pipette and electroporated (1500V 20ms 3 pulses) using the Neon transfection system device. After electroporation cells were immediately transferred in pre-warmed post-transfection medium (made with RPMI 1640 instead of *α*MEM and without antibiotics). Following 3h of incubation in the post-transfection medium, cells were pelleted by centrifugation, resuspended in fresh medium and grown in standard culture conditions.

After 16h, transfected cells were harvested and resuspended in RPMI 1640 with 2% FCS. Sorting was performed using BD FACS Aria 1 flow cytometer. Non-transfected cells were used as a negative control to discriminate between positive and negative GFP cells. Thus living GFP-expressing cells were collected, pelleted by centrifugation and grown back in LM1 or DM17 fresh media.

### ImageStreamX analysis

#### Sample preparation

Cells were pelleted by centrifugation, resuspended at the same concentration (2.10^6^ cells/ml). One part of the cells was incubated for 10 min with FCCP (carbonyl cyanide-p-trifluoromethoxyphenylhydrazone) (10 *μ*M) (ABCAM). FCCP was used as a negative control for mitochondrial membrane potential (MMP) measurements.

Cell were incubated 20 min in culture media completed with a staining solution composed of MitoTracker Green FM (100 nM) (BD Horizon), TMRE (ultra pure) mitochondria dye (Enzo) (1 nM) and Hoescht 33342 (In vitrogen) (2.5 *μ*g/ml). Following incubation cells were washed two times with 1X PBS. To discriminate between live and dead cells, they were finally incubated with Fixable Viability Stain 660 (FVS 660; BD Horizon) at 1:2000 dilution for 15 min at 4°C in the dark. Cells were washed once and resuspended in 1X PBS before measurements with ImageStreamX Mark II (ISX) (Amnis). A part of the cells were single stained in parallel as compensation samples.

#### Images acquisition

Stained cells were loaded within ISX device. During acquisition, cells were gated according to the area and aspect ratio features. For each condition, five thousand cells were recorded with 60x objective at the rate of 60 cell/s. Lasers 405, 488, 561, 642, 785 nm were used to analyze respectively DNA, mitochondria, mitochondrial membrane potential, dead cells, and cell granularity.

#### Image analysis

Data generated from ISX were analysed with the dedicated IDEAS software (Merck Millipore). First, image compensation was performed using single stained samples. Thereafter, cells were more precisely gated. Focused cells were gated first using bright-field and gradient RMS feature, from which singlets were gated using area and Hoechst intensity features, and finally living cells were gated from singlets, using intensity feature of FVS 660 (negative cells). Then, cellular mitochondrial content and MMP were assessed from green mitotracker intensity and TMRE intensity, respectively, using intensity features. Those data were then exported and analyzed with R software [18].

### Cell respiration measurements

Oxygen consumption rates were measured using Oroboros oxygraph (Oroboros Instruments), composed of two analysis chambers. Four respirometers were available and two conditions per apparatus could be tested, therefore eight conditions per experiment were performed. To avoid technical variability when comparing kinetic time points, conditions replicates where loaded randomly in different respirometers. One million cells per chamber were loaded at a final density of 5.10^5^ cells/ml in their culture media. During measurements cells were maintained at 37°C under constant stirring within the chambers.

Routine endogenous respiration was first measured after respiration stabilization in the chambers (Vendo), then oligomycin (1.25 *μ*g/mL), an inhibitor of the ATP synthase, was added and the non-phosphorylating respiration rate recorded (Voligo). Thereafter, a subsequent titration from 2.5 *μ*M to 10 *μ*M FCCP was performed to assess maximal electron transport chain activity (VFCCP). Antimycin A (22.5 *μ*M), an inhibitor of cytochrome bc1 complex (complex III of the respiratory chains), was added and the non-mitochondrial oxygen consumption rate of cells recorded (Vantimycin). Antimycin inhibited rates were equally low during differentiation process (20% ± 2% of routine), indicating that mitochondria accounted for 80% ± 2% of routine respiration. Finally, ascorbate was added at a final concentration of 2.5 mM followed by a subsequent titration of N,N,N’,N’-tetramethyl-p-phenylenediamine (TMPD) from 125 *μ*M to 750 *μ*M in order to measure the oxidative capacity of cells associated with isolated cytochrome-c oxidase activity. Using this approach, we determined the oxygen consumption associated with mitochondrial proton leak (Voligo - Vantimycin), mitochondrial ATP synthesis ([Vendo - Vantimycin] - [Voligo - Vantimycin]), and mitochondrial respiratory reserve ([VFCCP - Vantimycin] - [Vendo - Vantimycin]).

### Measurement of extra-cellular lactate concentration

Cells were grown during 2h in RPMI 1640 at the density of 2.10^6^ cells/ml and cellular supernatants were loaded in a 10 kDa ultracentrifugation filter (Amicon; Merk Millipore) to eliminate lactate dehydrogenase enzymes, as suggested by manufacturers protocol. Soluble fractions were then assessed using the Lactate assay kit (SIGMA) according to manufacturers procedure, and samples absorbance was measured at 570 nm using the Multiskan GO plate reader (Thermo Scientific). Background absorbance was substracted from samples reading and lactate amount was determined using a standard dilution curve supplied by the Lactate assay kit. Lactate concentration was then calculated as recommended by manufacturers protocol.

### Proteomic data generation

Proteomic data were generated using a label-free quantification (LFQ) method ans mass spectrometry by the Plateforme de Protéomique de l’Université Paris Descartes (3P5) [19].

### CRISPR plasmid construction

#### T2A substitution by P2A in PX458 vector

It has been shown in various species that P2A (2A peptide derived from porcine teschovirus-1) presents a higher cleavage efficiency than T2A (2A peptide derived from Thoseaasigna virus) [20], which was initially present in pSpCas9(BB)-2A-GFP (PX458) vector. pSpCas9(BB)-2A-GFP (PX458) was a gift from Feng Zhang (Addgene plasmid # 48138; http://n2t.net/addgene:48138; RRID:Addgene 48138) [21]. Thus we decided to substitute T2A for P2A to enhance vector efficiency.

Oligonucleotides encoding P2A and homologous sequences were purchased from Eurogentec (table 1). P2A was substituted for T2A using the PCR Geiser method [22]. P2A oligonucleotides were used as mega primers for PCR on PX458 vector.

**Table 1.**
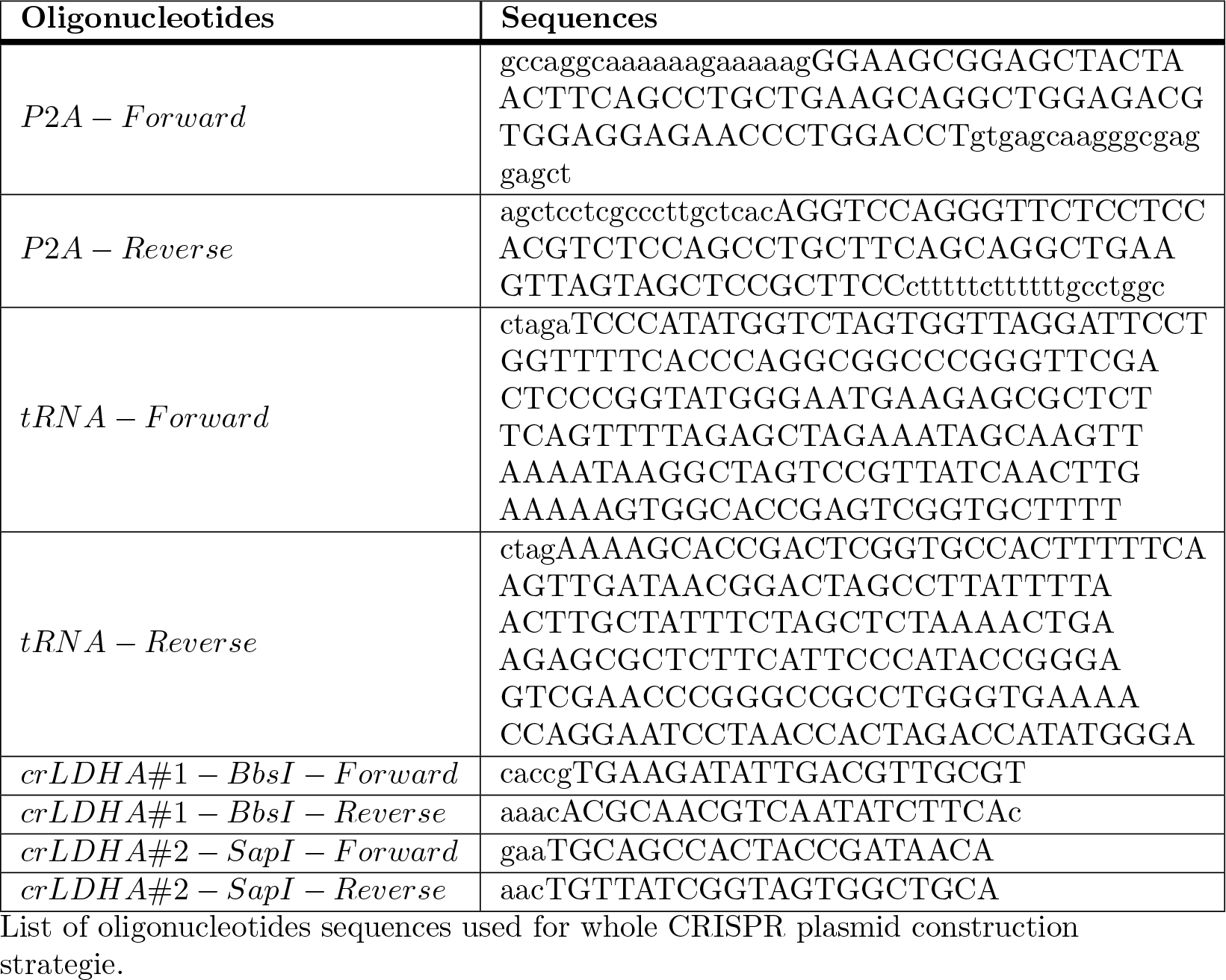
Oligonucleotides sequences used for CRISPR plasmid construction.

#### Multiplex CRISPR system construction

pCRISPR-P2A vector was modified to produce a multiplex CRISPR/Cas9 system to allow expression of two RNA guides (gRNA) with the same vector. Therefore we added a tRNA and a new restriction multiple site to insert a second gRNA in pCRISPR-P2A vector. Oligonucleotides encoding chicken tRNA Glu and RNA scaffold were purchased from Eurogentec, annealed and then cloned into XbaI-digested pCRISPR-P2A vector. After validation by sequencing the final plasmid was named pCRISPR-P2A-tRNA.

#### LDHA gRNA cloning

Two RNA guides (crLDHA#1 and crLDHA#2) against LDHA sequence were designed using CRISPR design tool (http://crispr.mit.edu) to target the first exons of the sequence. The first guide was cloned after hU6 promoter into BbsI-digested pCRISPR-P2A-tRNA vector. The second guide (crLDHA#2) was cloned after chicken tRNA Glu into SapI-digested pCRISPR-P2A-tRNA vector.

The efficiency of our CRISPR vector was confirmed by analyzing LDHA deletion by PCR on transfected cells (data not shown).

### Quantitative Real-Time Polymerase Chain Reaction assays

Total RNA was extracted from the transfected cells using the RNeasy Mini kit (Qiagen) according to manufacturer’s protocol. Reverse transcription was performed using Invitrogen™ SuperScript™ III First-Strand Synthesis SuperMix for qRT-PCR (Thermo Fisher Scientific) according to manufacturer’s protocol. q-PCR reactions were performed with SYBR Premix Ex Taq (TaKaRa) and analyzed using CFX connect Real-Time PCR detection system (Bio-Rad). Primers for the qPCR were purchased from Euroventec (table 2).

**Table 2.**
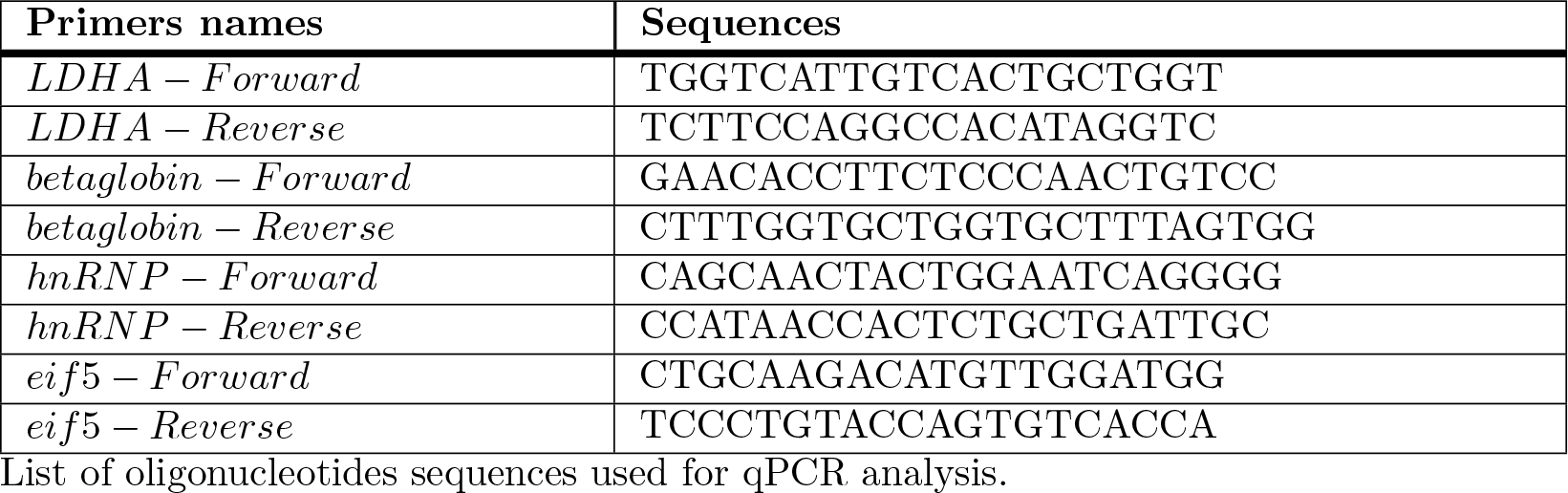
Primers sequences used for q-PCR analysis.

### Statistical analysis

Every statistical analyses were performed using R software [18]. The normality of each distribution was checked, using either qqplots or the shapiro test, in order to compute the appropriate tests.

## Results

### Variations of the level of glycolytic enzymes during erythroid differentiation

We first analyzed protein levels of glycolytic enzymes, in self-renewing and in differentiating primary chicken erythroid progenitor cells (T2EC). In order to dissect energetic glycolytic metabolism we selected important proteins from the following categories : (1) proteins involved in glycolysis, (2) proteins involved in oxidative phosphorylation (OXPHOS), and (3) the LDHA enzyme, involved in the last reaction of glycolysis, catalyzing lactate production (Fig 1A). Glycolysis is finely regulated at three key rate-limiting steps catalysed by Hexokinases (HK), Phosphofructokinase (PFKP) and Pyruvate kinase (PKM) [23–25]. Thus we decided to analyse the expression of these enzymes including two isoforms, HK1 and HK2, of hexokinase (Fig 1B). Globally HK1, PFKP and PKM expression was sharply decreased (more than 70%) during the differentiation process, whereas HK2 expression decreased to a lesser extend (less than 50%). However PKM protein level increased slightly (around 10%) at 12h. The important decline of the expression level of these proteins suggested that self-renewing T2EC displayed a high glycolytic flow, that was strongly reduced during differentiation.

**Fig 1.**
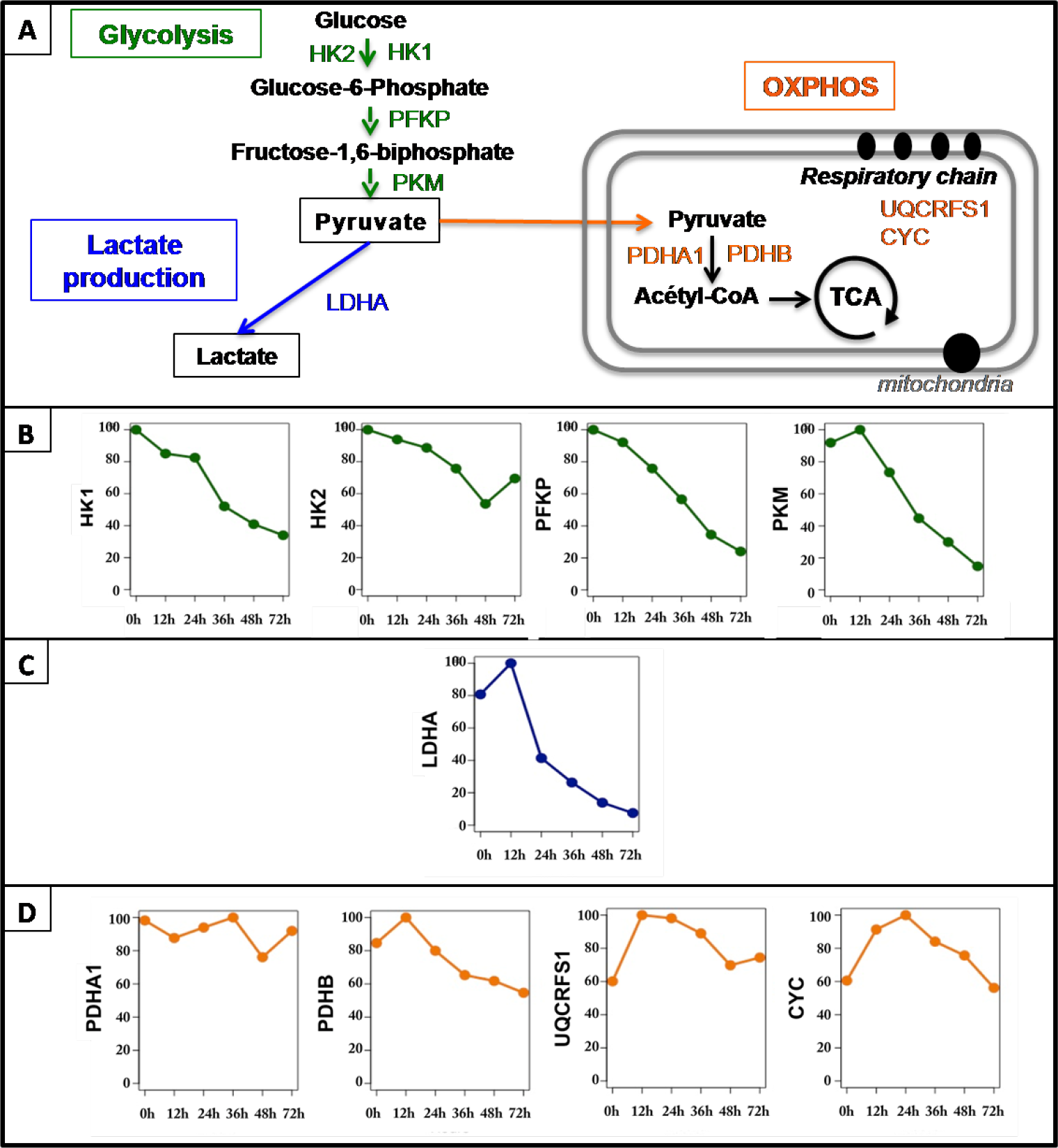
Relevant proteins for glycolytic metabolism and OXPHOS analysis. A: Proteins were selected from three different categories : key enzymes involved in glycolysis regulation (Glycolysis), LDHA alone, and OXPHOS-related enzymes (OXPHOS). B-D: Protein quantification was performed using mass spectrometry (MS/MS) upon three independent self-renewing T2EC populations and three independent T2EC differentiation kinetics (T2EC induced to differentiate for 12, 24, 36, 48 and 72h). Mean protein levels were normalized as follows : for each enzyme the maximal value was considered equal to 100, and all other values were proportionally converted. Normalized protein levels are represented on the y-axis whereas x-axis correspond to differentiation kinetic time-points (hours). LDHA: Lactate dehydrogenase A, HK1: Hexokinase isoform 1, HK2: Hexokinase isoform 2, PFKP : Phosphofructokinase Platelet, PKM: Pyruvate Kinase Muscle, PDHA1: Pyruvate Dehydrogenase E1 Alpha 1 Subunit, PDHB: Pyruvate Dehydrogenase E1 Beta Subunit, UQCRFS1: Ubiquinol-Cytochrome C Reductase, Rieske Iron-Sulfur Polypeptide 1, CYC: Cytochrome C.

We also analyzed the protein level of the enzyme involved in the last reaction of glycolysis involving lactate production, LDHA (Fig 1C). LDHA expression first increased (20%) at 12h, and then dropped massively (more than 80%) until 72h. Such an important decrease of LDHA protein expression implies a decrease of pyruvate conversion into lactate and suggest that glycolysis might be reduced during erythroid differentiation.

Pyruvate resulting from glycolysis can either be converted to lactate, or be transported into mitochondria to participate to OXPHOS, to produce ATP. Briefly, into the mitochondrial matrix pyruvate must be converted to acetyl CoA and enter in the tricarboxylic acid (TCA) cycle to generate electron transporters, NADH,H+ and FADH2, in order to feed respiratory chains coupled with ATP production, i.e. OXPHOS (Fig 1A). Thus, we included in OXPHOS genes category, pyruvate dehydrogenase complex subunits, PDHA1 and PDHB, that catalyse pyruvate conversion into acetyl CoA [26, 27]. Moreover we also selected the ubiquinol-cytochrome c reductase (UQCRFS1), a key component of complex III of the respiratory chains [29] and cytochrome c electron carrier (CYC) involved in OXPHOS [28] (Fig 1D). PDHA1 expression do not show any clear tendency during differentiation despite the general gene expression decrease, suggesting that it is particularly needed. However, PDHB expression first increased slightly (20%) at 12h and then declined slowly. Although PDH subunits level gave interesting information, it is important to note that PDH complex activity, as measured by its phosphorylation level, was not evaluated in this study. Expression of respiratory chains components UQCRFS1 and CYC displayed an interesting profile. They first show a 40% raise at 12-24h of differentiation and then returned progressively to the protein level displayed in self-renewing T2EC. Increased expression of PDH complex subunit PDHB might suggest that more pyruvate molecules could be recruited in the mitochondria to be converted into acetyl CoA at 12h of the differentiation process. Moreover the increase of UQCRFS1 and CYC proteins also suggests that respiratory chains activity might be boosted at that stage of the differentiation process. It is important to note here that during erythroid differentiation, gene expression decreases progressively because of the nucleus condensation [15]. The slight decrease of the respiratory chains proteins after 24h thus, could be due to this general decline, which does not necessarily mean that OXPHOS capacity is reduced.

Overall, during T2EC differentiation, LDHA and PKM protein levels increased at 12h, and then decreased significantly. The other glycolysis-related enzymes decreased gradually from the self-renewal state (0h) up to 72h. OXPHOS-related enzymes, including PDHB subunit of the PDC complex, increased at 12-24h, whereas PDHA1 subunit remained most likely stable over differentiation. One can therefore suggest that erythroid differentiation might be accompanied by an enhancement of the respiratory chains and glycolysis activities specifically at 12h, followed by a strong decline of the glycolytic pathway and back to a normal level of OXPHOS.

### Energetic glycolytic metabolism changes during erythroid differentiation

We first compared the capacity of self-renewing and differentiating T2EC to produce lactate. We observed that the differentiation was accompanied by a significant decrease in the ability of our cells to produce lactate (Fig 2). This result was in line with observed decline of LDHA protein expression (Fig 1), suggesting that lactate producing glycolysis was indeed reduced during the differentiation process. We then wanted to evaluate whether glycolytic metabolism resulting in lactate production might switch toward OXPHOS during the erythroid differentiation process. As a measure of mitochondrial activity, we decided to use TMRE, a fluorescent dye which accumulates within mitochondria according to the mitochondrial membrane potential (MMP), reflecting proton motive force. However, since cell size strongly decreases during differentiation [13] and that the overall amount of mitochondria has been previously shown to correlate with cell size [30], we first assessed the overall amount of mitochondria using a green mitotracker. Mitotracker intensity dropped at 24h of the differentiation process and remained low until 72h (Fig 3). Given that mitochondria density declined strongly at 24h, we considered that effect in our next mitochondria-dependent measurements.

**Fig 2.**
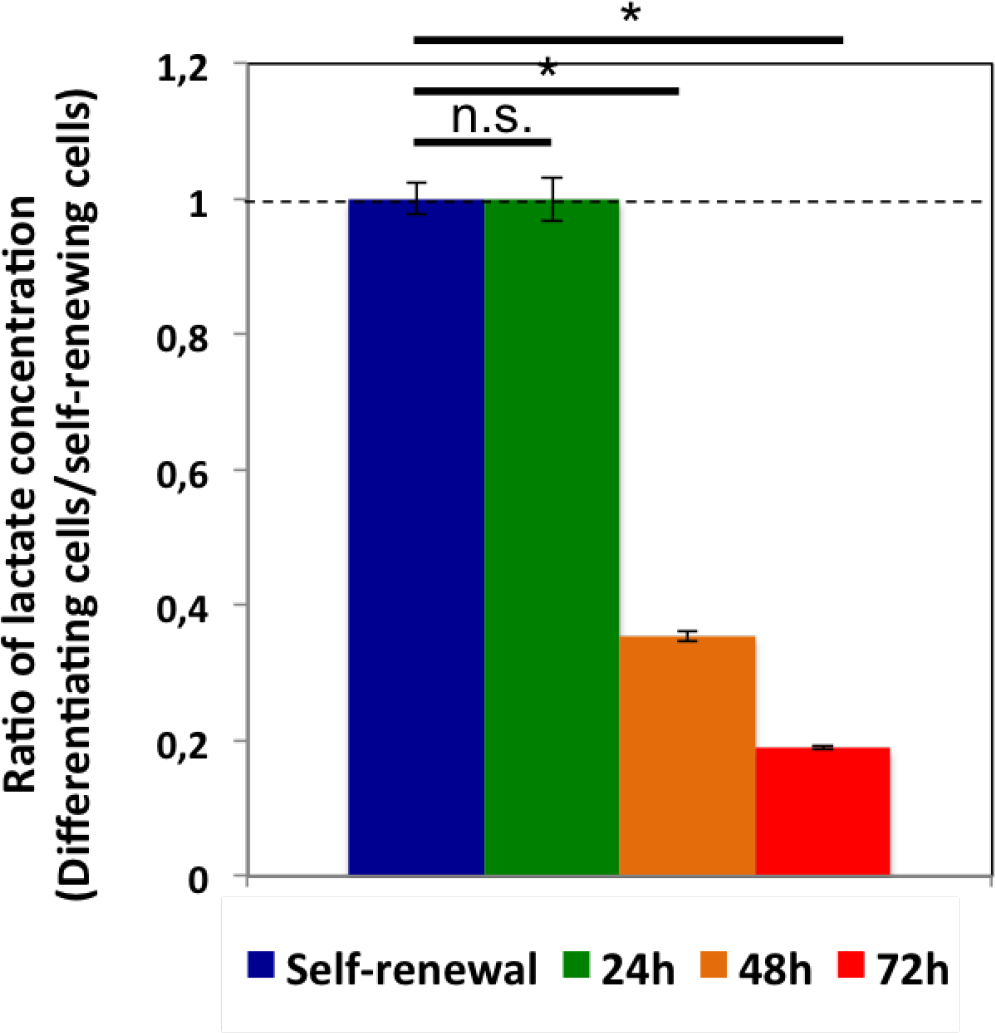
Lactate production by self-renewing and differentiating cells. Extra-cellular lactate concentration in self-renewing cells and cells induced to differentiate for 24h, 48h and 72h was assessed following incubation during 2h in fresh media. Represented is the ratio of extra-cellular lactate concentration of each condition divided by the self-renewal condition. Lactate concentration of self-renewing cells is equal to 1, as divided by itself, and indicated by the dotted line. Bars represent mean +/− S.D. calculated from four independent experiments for 24h and 48h differentiation time-points and five independent experiments for 72h and self-renewal. A t-test was applied to assess whether means were significantly different (*: p-value < 0.05; n.s.: non significant).

**Fig 3.**
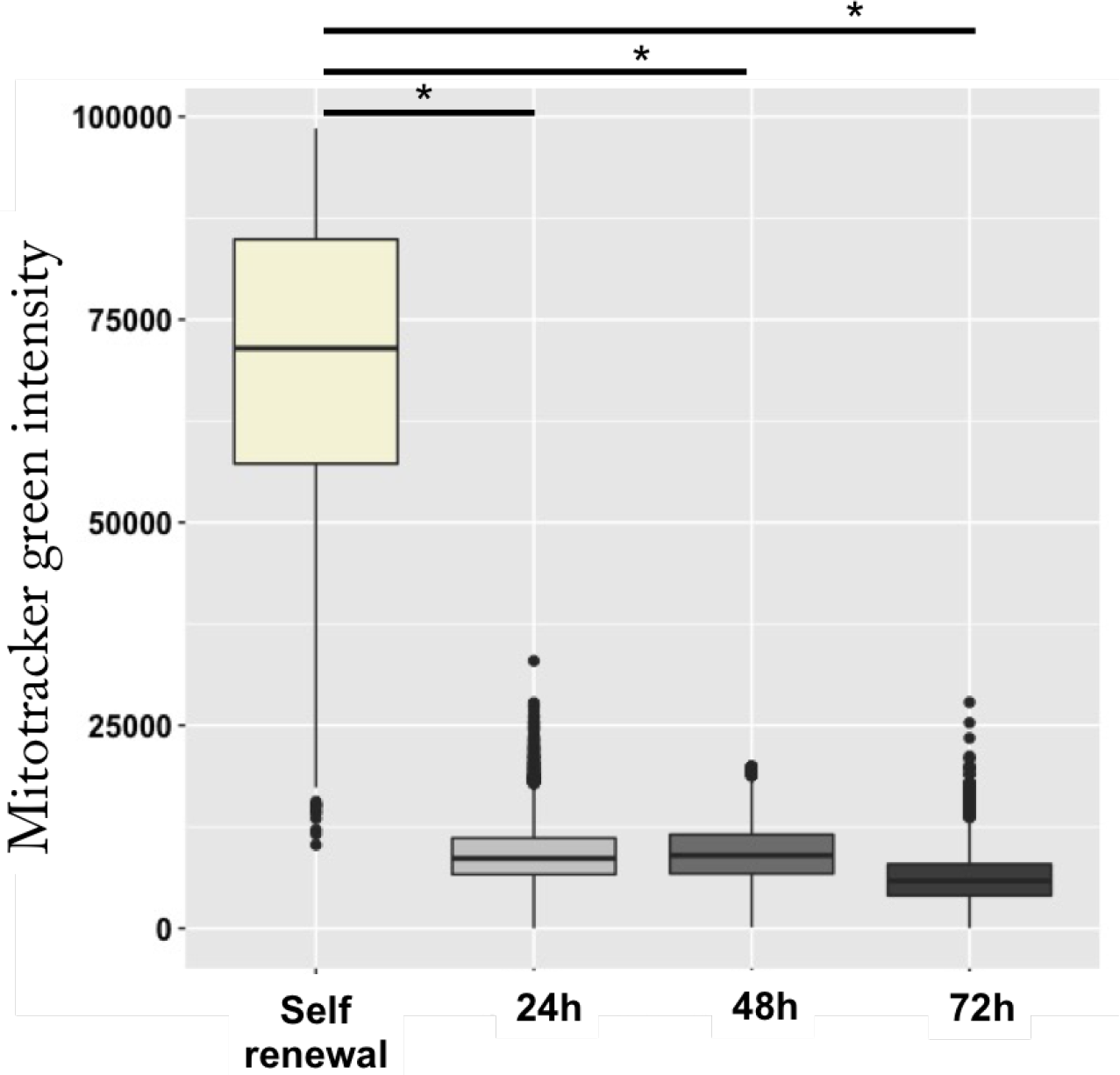
Mitotracker intensity evolution during the erythroid differentiation process. Shown is the distribution of the mitotracker intensity at each time-point of the differentiation process measured using ImageStreamX. A t-test was applied to assess whether means were significantly different (*: p-value < 0.05). Data were obtained from three independent experiments. 11678 cells were analyzed for the self renewal condition, 12587 cells for 24h, 14147 cells for 48h and 12355 cells for 72h.

When the mitochondrial membrane potential (MMP) is dissipated by uncoupling molecules, such as FCCP, TMRE does not accumulate anymore within mitochondria. Thus, we used the FCCP uncoupler as a control for TMRE staining. In the presence of FCCP, only TMRE staining disappeared, reflecting the specific response of TMRE to the dissipation of MMP following mitochondrial uncoupling, and mitotracker independence from MMP. It also confirmed that the TMRE concentration used is a non-quenching dose (Fig 4A). In order to analyze only living cells, we also used FVS 660 dye, that accumulates within dead cells (Fig 4B). Therefore FVS 660 positive cells were removed from our analysis.

**Fig 4.**
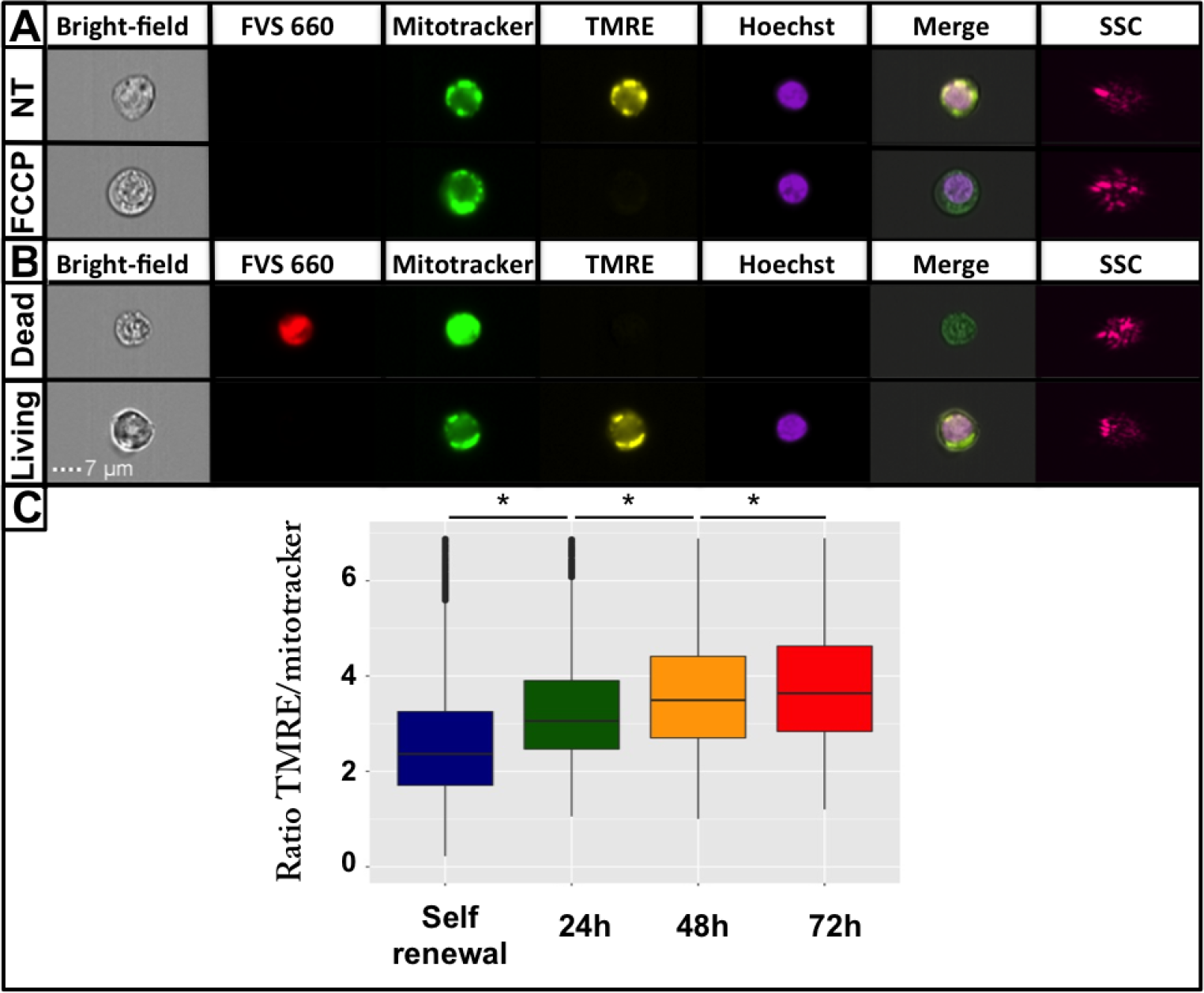
Mitochondrial membrane potential (MMP) evolution during erythroid differentiation. Cells morphology and size were observed through the bright field channel using ImageStreamX. Dead cells were identified using FVS 660 dye (red). MMP was assessed using TMRE dye (yellow), mitochondrial content was evaluated using green mitotracker intensity (green), and the nucleus was stained using Hoechst (purple). Merge corresponds to the superposition of bright field, TMRE, mitotracker and Hoechst images. Cell granularity was assessed using SSC channel (pink). A: The FCCP uncoupler was used as a negative control of MMP staining. NT: No treatment. B: Shown is an example of FVS 660 staining to discriminate between living and dead cells. C: Boxplots of normalized TMRE distributions. Outliers are not shown. A Wilcoxon test was applied to assess whether means were significantly different (*: p-value < 0.05). Data were obtained from three independent experiments. The overall numbers of cells analyzed were 11678 for the self-renewal condition, 12587 cells for 24h, 14147 cells for 48h and 12355 cells for 72h.

Finally, to account for cellular mitochondrial content variations, we normalized the fluorescent TMRE intensity with the green mitotracker intensity. We then assumed that the MMP corresponds to the ratio of TMRE upon mitotracker intensity. Consequently, we plotted this ratio and analyzed MMP behavior during the differentiation process (Fig 4C). We noted that MMP increased significantly during erythroid differentiation. Such an increase was a strong indication of an enhanced OXPHOS, and thus tend to support the hypothesis that T2EC switched from glycolysis toward OXPHOS during their differentiation process. In order to study OXPHOS activity into finer details, we assessed oxygen consumption in self-renewing and differentiating T2EC (Fig 5). We first observed that routine cell respiration was significantly increased in T2EC induced to differentiate for 24h, and then decreased at 48h and 72h to reach a similar level than self-renewing T2EC respiration rate (Fig 5A). Different calculations were then performed to better decipher the respiration mechanisms (see subsection *Cell respiration measurements* for more details). First, we evaluated oxygen consumption resulting from proton leak. The oxygen consumption associated with mitochondrial proton leak activity increased significantly at 24h and decreased significantly at 48h (data not shown). Moreover, proton leak activity represented on average 34% ± 3% of routine respiration. In turn this implies that 66% ± 3% of oxygen consumed by mitochondria was dedicated to synthesize ATP, regardless of the differentiation stage (Fig 5B). Similarly to routine respiration profile along differentiation, the oxygen consumption devoted to ATP production increased at 24h and then decreased significantly to come back to the initial respiration rate at 72h (Fig 5B). Therefore, the surge observed in routine respiration at 24h might be due to an increase of cellular energy needs. Moreover, as shown upstream, cellular mitochondrial content decreased during the differentiation process as soon as 24h (Fig 3) suggesting that the increased cellular respiration at that stage did not rely upon an increase in mitochondrial content, but mostly on an increase of the mitochondrial OXPHOS activity.

**Fig 5.**
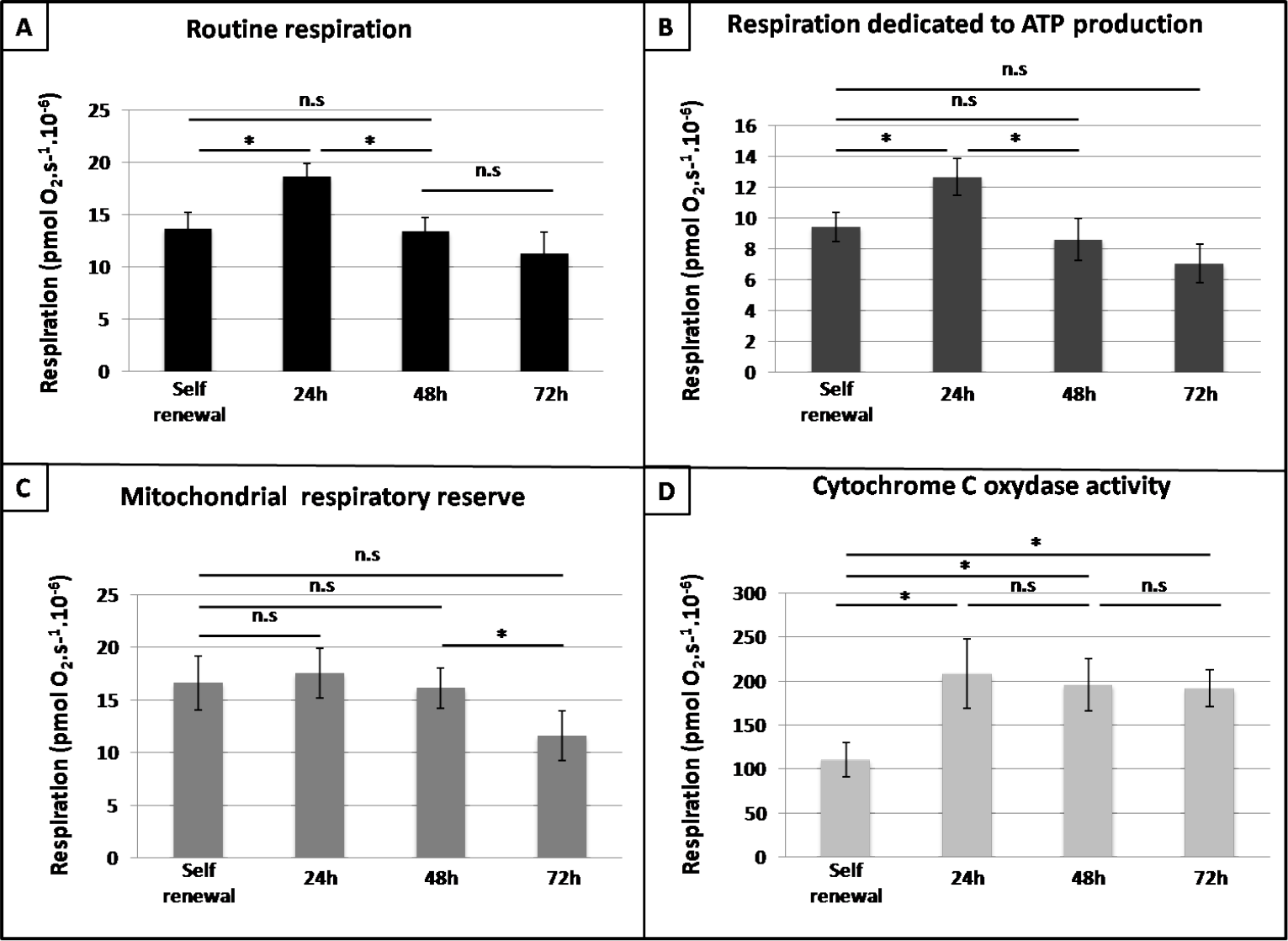
Dissection of cellular respiration during the differentiation process. Different respiration parameters were assessed in self-renewing T2EC and T2EC induced to differentiate for 24, 48 and 72h. A: Routine respiration corresponds to oxygen consumption without any addition. B: Respiration dedicated to ATP production was evaluated by subtracting respiration after oligomycin treatment from routine respiration. C: Subtraction of routine respiration from maximal uncoupled respiration by FCCP reflected mitochondrial respiratory reserve. D: Cytochrome C oxydase activity was assessed by subtracting respiration following ascorbate addition from maximal TMPD-related respiration. Bars represent means +/− S.E.M. from five independent experiments performed in duplicates. A paired t-test was applied to assess whether means were significantly different (*: p-value < 0.05; n.s.: non significant).

We also assessed maximal respiration capacity by subtracting routine respiration from the respiratory reserve capacity obtained by uncoupling OXPHOS with FCCP (Fig 5C). We observed no significant variations of respiration capacity except a decrease between 48h and 72h.

Finally, activity of cytochrome C oxidase, reflecting the oxidative capacity of cells, was estimated (Fig 5D). Cell oxidative capacity was significantly increased at 24h and remained stable until 72h, suggesting that differentiating T2EC displayed a better capacity to perform OXPHOS than self-renewing T2EC, confirming MMP measurements.

Taken together these results strongly suggested that during erythroid differentiation, energetic demand increased at 24h, which was satisfied by a boost of OXPHOS, and that cell metabolism switches from glycolysis toward OXPHOS during erythroid differentiation.

### LDHA might be involved in glycolytic changes during erythroid differentiation

We showed that LDHA protein expression decreased strongly during the differentiation process (Fig 1C). This suggests that a reduction of LDHA activity might be necessary for cells to commit to differentiation. Furthermore, since LDHA is the most important enzyme of the glycolytic pathway characterized by lactate production, the necessary decline of its expression might be involved in the metabolic switch. To assess whether LDHA activity could influence the metabolic status of T2EC, we inhibited LDHA activity using Galloflavin and FX11, two known LDHA inhibitors [31–33], and measured the MMP.

We first determined non-cytotoxic concentrations of Galloflavin (50 *μ*M) and FX11 (30 *μ*M) by titration (data not shown). We then assayed Galloflavin and FX11 efficiency by measuring extra-cellular lactate concentration after treating T2EC for 24h with these inhibitors. As expected, lactate production was diminished following LDHA inhibitors treatments (Fig 6A). We then incubated T2EC with Galloflavin or FX11, for 24h and measured MMP using ISX (Fig 6B). Following inhibitor treatment MMP increased significantly, mostly after FX11 treatment, that has the most important effect on lactate production. Consequently, the decrease of LDHA activity was accompanied with an increase of mitochondrial activity, suggesting that LDHA might indeed be involved in metabolic status changes. These results highlighted the link between the rapid drop of LDHA protein level and the switch toward OXPHOS occuring during the erythroid differentiation process.

**Fig 6.**
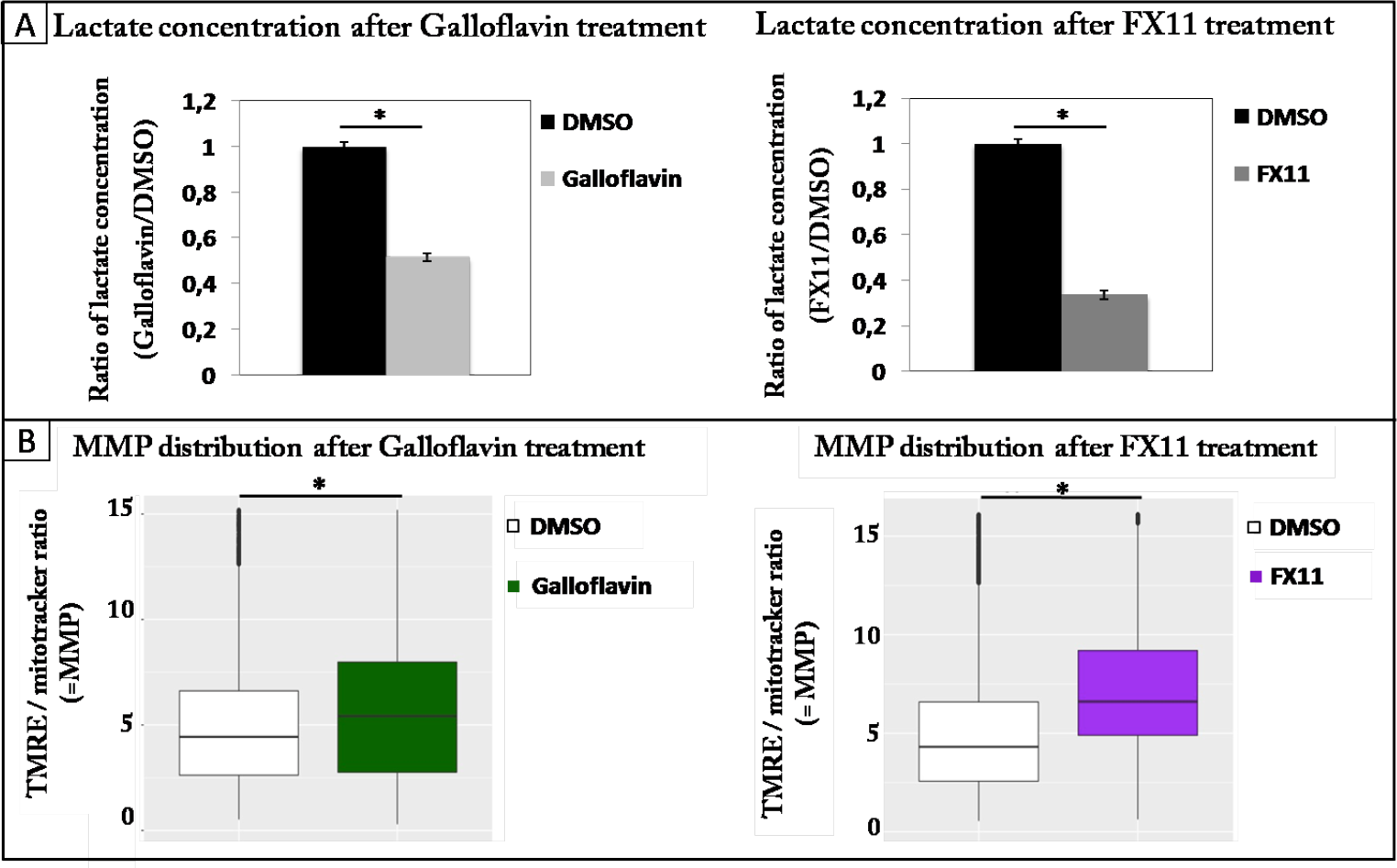
Mitochondrial membrane potential changes following LDHA inhibition. T2EC were incubated with Galloflavin (50 *μ*M) and FX11 (30 *μ*M) for 24h. A: Lactate concentration was measured in T2EC media following incubation with Galloflavin or FX11 for 24h. Each value represents mean +/− S.D. of six independent experiments for the Galloflavin and five independent experiments for FX11. A t-test was applied to assess whether distributions were significantly different (p-value inf. to 0.05). B: MMP was assessed on living cells (FVS 660 dye) using TMRE dye, normalized by dividing TMRE intensity by mitotracker intensity. FCCP uncoupler was used as a negative control for MMP staining. A wilcoxon-test was applied to assess whether distributions were significantly different (p-value inf. to 0.05). Data were obtained from three independant experiments. The overall numbers of cells analyzed for Galloflavin treatment were respectively 11306 and 11994 cells for the DMSO and Galloflavin conditions. The overall numbers of cells analyzed for FX11 treatment were respectively 10904 and 12858 cells for the DMSO and FX11 conditions.

### Maintenance of erythroid progenitors self-renewal required LDHA expression

The important decrease of LDHA expression during the erythroid differentiation process suggested that LDHA might be important to maintain self-renewal of T2EC. We therefore inhibited LDHA activity in self-renewing T2EC (Fig 7A). Either FX11 or Galloflavin treatment slowed down significantly cell growth, suggesting that LDHA activity is important for T2EC self-renewal. To confirm these results we used the CRISPR-cas9 technology to inhibit *LDHA* expression, and evaluated cell growth and differentiation rate. As observed with Galloflavin and FX11 treatments, the inhibition of *LDHA* expression with the CRISPR vector, tended towards slow down cell growth compared to cells transfected with an empty vector (Fig 7B). We therefore confirmed the involvement of LDHA in the self-renewal of erythroid progenitors.

**Fig 7.**
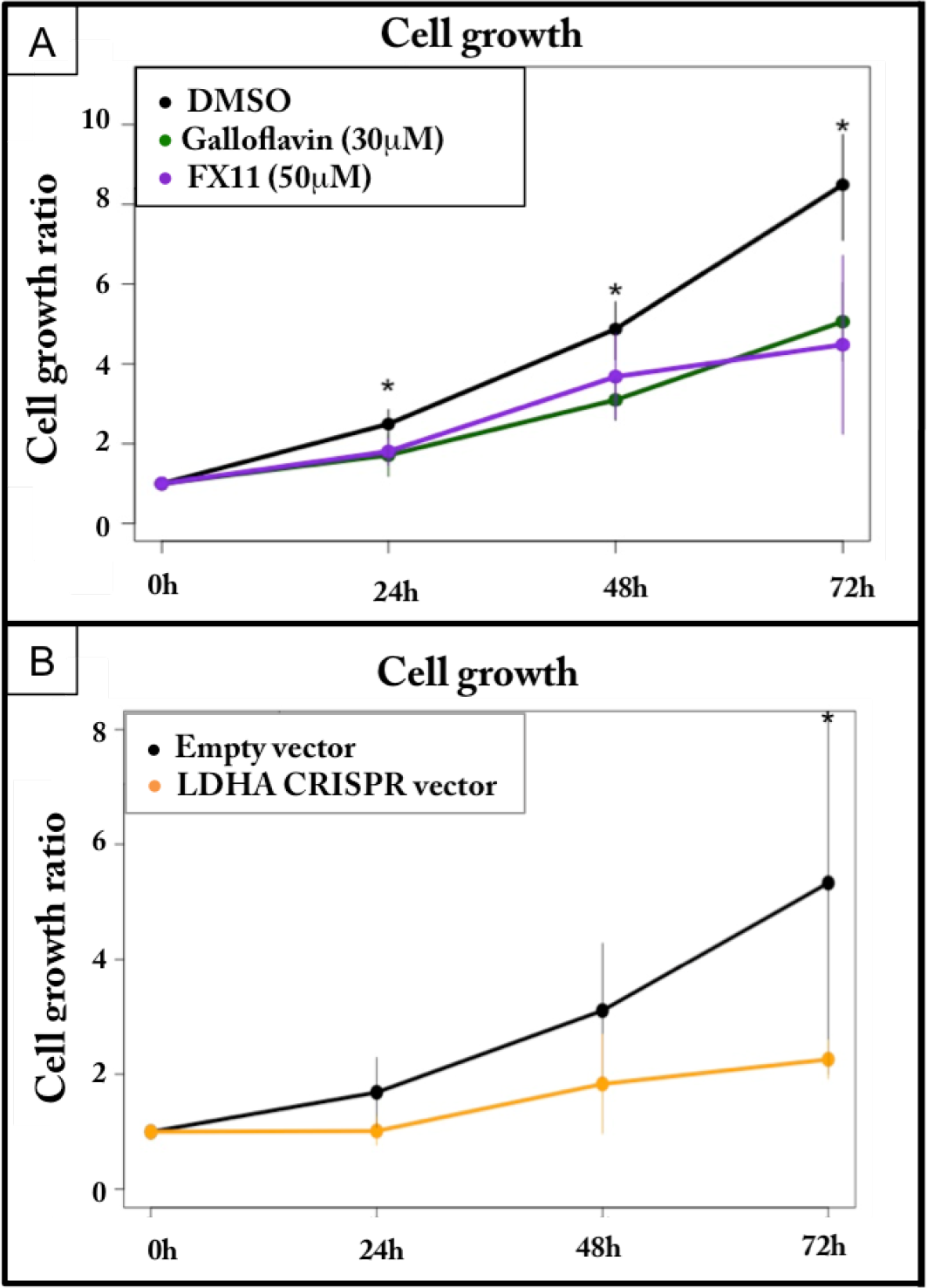
Consequences of LDHA inhibition on the growth of self-renewing cells. A: Self-renewing T2EC were incubated with Galloflavin or FX11 and were counted every 24 hours for 3 days. Results are the mean +/− S.D. calculated from four independent experiments for FX11, and five independent experiments for Galloflavin. Growth ratios were calculated as the cell number divided by the total cells at day 0. The significance of the difference (p-value inf. to 0.05) between control growth ratios and LDHA inhibitors were calculated using a Wilcoxon test. B: T2EC were transfected with a CRISPR vector directed against *LDHA* or an empty vector, and were counted every 24 hours for 3 days. The orange curve indicates the growth of T2EC with the CRISPR vector and the black curve indicates the growth of T2EC with the empty vector. The data shown are the mean +/− S.D. calculated from three independent experiments. Growth ratios were calculated as the cell number divided by the total cells at day 0. The significance of the difference (p-value inf. to 0.05) between control growth ratios and LDHA CRISPR were calculated using a t test.

Otherwise, the inhibition of *LDHA* expression, using CRISPR technology and inhibitors, did not affect T2EC differentiation, which was consistent with the fact that LDHA expression is already sharply decreased under such condition (data not shown). Since the inhibition of LDHA activity or quantity did not affect the differentiation process, it suggests that the slight increase of its protein expression at 12h (Fig 1C) is not essential for cells to engage into differentiation.

### The metabolic switch toward OXPHOS might be a driving force for erythroid differentiation

Our results suggested that a metabolic switch might be occurring during the erythroid differentiation process. However whether or not these metabolic rearrangements could be involved in T2EC differentiation was still unknown.

To assess the role of such metabolic shift on T2EC ability to differentiate, OXPHOS was impaired during the differentiation process, using the respiratory chains complex III inhibitor, antymicin A. T2EC were induced to differentiate in presence of antimycin A or DMSO during 48h and cell differentiation rate was assessed following two strategies. We first used benzidine to compare the percentage of differentiating cells between both conditions (Fig 8A). Our results showed that antimycin A treatment significantly impaired T2EC differentiation. We then quantified the relative expression of *betaglobin*, a marker of erythroid differentiation, and *LDHA* genes (Fig 8B). *LDHA* expression was increased whereas *betaglobin* expression was decreased following antimycin A treatement, in respect to DMSO control condition. Consequently, OXPHOS impairment seemed to affect negatively the differentiation process.

**Fig 8.**
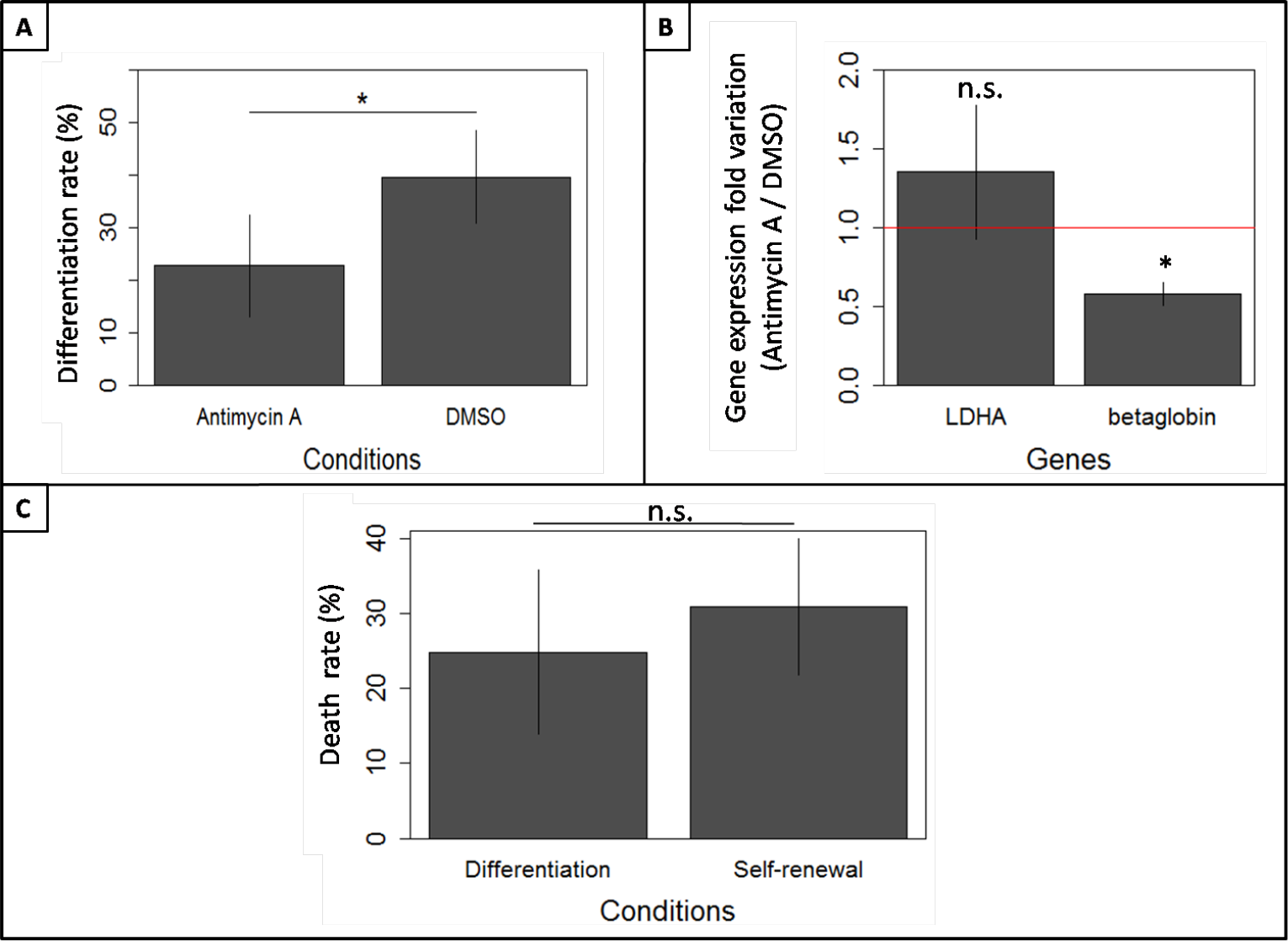
Consequences of OXPHOS impairment on T2EC differentiation. A: T2EC were incubated in differentiation media with antimycin A (10nM) or DMSO, during 48h. The differentiation rate was assessed using benzidine. The percentage of differentiating cells was calculated as the benzidine positive cells divided by the total cells. Results are the mean +/− S.D. from four independent experiments. The significance of the difference (p-value inf. to 0.05) between control and antimycin A-treated cells was calculated using a t-test. B: LDHA and betaglobin relative mRNA levels were quantified in T2EC treated with either antimycin A (10nM) or DMSO, during 48h in differentiation media. Data were normalized according to *hnRNP* and *eif5* standard genes. The data shown are the mean fold changes +/− the standard deviation of gene expression under Antimycin A treatment with respect to the control (DMSO) condition, from three independent experiments. The red line represents fold change value y=1. C: Mortality rate was compared between T2EC treated with antimycin A (10nM) in self-renewal (normal culture conditions) and differentiation media, for 48h. Mortality ratios were calculated as the trypan positive cell number over total cell number. The data shown are the mean +/− S.D. calculated from four independent experiments. An unilateral t-test was applied to assess whether means were significantly different (*: p-value < 0.05).

However, when analyzing gene expression, only living cells were harvested and therefore considered. The observed effect could then either result from: (1) a higher sensitivity of differentiating cells to antimycin A treatment, which would therefore die more than self-renewing cells, or (2) a *bona fide* block in differentiation induced by antimycin A treatment. To decide between those two possibilities, we compared self-renewing and differentiating T2EC sensibility to antimycin A treatment (Fig 8C). Our results clearly show that self-renewing T2EC are as sensitive as differentiating T2EC to Antimycin A treatment. Consequently, variations of *LDHA* and *betaglobin* expressions, following antimycin A treatment, did not result from an increased mortality rate of differentiating cells, but rather from an intrinsic decrease of T2EC capacity to differentiate.

Therefore, our results showed that OXPHOS impairment affected the differentiation process, suggesting that the switch from glycolysis involving lactate production toward OXPHOS might play an important role in erythroid differentiation.

## Discussion

This study aimed to determine how glycolytic metabolism changes during the erythroid differentiation process, and which role such metabolic rearrangements might play in erythroid progenitors metabolic status. In this context we used a multi-parametric approach including molecular analysis, metabolic parameter measures and cellular physiological insights.

To obtain a glimpse of the changes that might occur in the glycolytic metabolism during erythroid differentiation, we defined a subset of nine enzymes, involved either in glycolytic flow regulation, OXPHOS or lactate production. The protein profile of these enzymes suggested that glycolysis flow and lactate production were increased at 12h and then slowed down until 72h. Therefore, besides the increase at 12h, glycolysis seemed to be impaired during erythroid differentiation. On the other hand, respiratory chains activity, as measured by the protein level of respiratory chains complex subunits, seemed to increase at 12h-24h, before returning to its initial level. Lactate concentration and mitochondrial membrane potential (MMP) assessments reinforced our molecular analysis, demonstrating that self-renewing progenitors produced more lactate than differentiating cells, and that MMP increased progressively during differentiation. Furthermore, we showed that LDHA inhibition increased MMP and slowed down cell growth. Concomitant with this result, cell respiration rate peaked at 24h, mostly due to ATP production, and most importantly, cells oxidative capacity was intensified in differentiating cells. Finally, OXPHOS inhibition impaired erythroid differentiation as assayed using antimycin A respiratory chains inhibitor.

All these results suggested that (i) self-renewing progenitors rely upon glycolysis through LDHA, (ii) a surge in energy demand occurs during the first phase of differentiation, and (iii) erythroid differentiation is accompanied by a metabolic switch from lactate-producing glycolysis toward OXPHOS, that could be necessary for the differentiation process.

### There is more metabolism than just glucose metabolism

In the present work we focused on glucose metabolism, from which LDHA is a key actor. LDHA is important to maintain glycolysis, specially when acetyl CoA level and TCA activity are low, as it regenerates NAD+ cofactors, required to run glycolysis. The simultaneous co-inhibition of LDHA and HK2 can induce the death of cancer cells that depend upon glycolysis [34]. However, the inhibition of LDHA only rather impair cell growth and tumorigenesis in the context of cancer [35, 36]. Similarly LDHA inhibition in human progenitor cells reduces cell growth, in accordance with our findings [2].

However, the glycolytic pathway is part of a broader network where glucose metabolism is highly connected with lipid and amino acid metabolisms. Interstingly, glutamine metabolism was shown to participate to erythroid differentiation [37]. Sterol metabolism has also been highlighted in erythroid cells. Indeed, it was demonstrated in our team, that Oxydosqualene Cyclase (OSC), which is involve in cholesterol biosynthesis, is needed to maintain the self-renewal state of erythroid progenitors [38]. We have also observed that sterol-related genes were early drivers of the differentiation process, as their expression changed as soon as 2h after the induction of differentiation [13]. Therefore, the orchestration of glucose and sterol metabolisms during erythroid differentiation could be a promising topic to study, that would bring new insights regarding metabolism implication in erythroid differentiation.

The generation of time-stamped metabolomic data along differentiation should be useful to acquire a more global point of view of the metabolic network and better explain energetic fluctuation and its repercussions, particularly regarding OXPHOS, the point of convergence of many metabolic pathways.

### May glucose metabolism switch drive erythroid differentiation?

There is increasing evidence from the literature that cells could adapt their metabolism in accordance with their environment and energy needs [39]. Hematopoietic progenitor cells and long-term hematopoietic stem cells, for instance, are dependent upon glycolysis and switch toward mitochondrial oxidation while they differentiate [40, 41]. On the other hand mature naïve T cells, use OXPHOS to produce energy, and shift back to the glycolytic pathway once they are activated [42]. Monocytes are also able to switch from glycolysis toward OXPHOS, and to switch back then to the glycolytic pathway, when they faced a microorganism [39].

A relevant review [8] suggested that a switch from glycolysis to OXPHOS might happen during erythroid differentiation. Such a view is fully supported by our results. Moreover, it is also interesting to note that a recent proteomic analysis performed on human erythroid differentiation, showed expression profiles for glucose metabolism-related enzymes (HK1, PFKP, PKM and LDHA) that are very similar to those observed in our avian erythroid cells [15].

One key question is then to what extent this metabolism change might be necessary for erythroid cells to differentiate, and if it might drive cells into the differentiation process, as it has been proposed for other cell types [4, 43–45]. Our finding regarding erythroid cells also support this hypothesis. The impairment of OXPHOS during erythroid differentiation using antimycin A affected cells ability to differentiate. Moreover, under these conditions LDHA expression tended to increase. We have shown that this was not due to a higher sensitivity of self-renewing cells to OXPHOS breakage. We propose that cells confronted with differentiation signals, but deprived of the appropriate metabolic machinery (OXPHOS), might adapt their metabolism by increasing glycolysis and pyruvate conversion into lactate to produce energy, which prevent them from leaving the self-renewal state.

Besides the global protein level decline of enzymes involved in glycolysis and pyruvate conversion into lactate, our analysis highlighted that some of these key enzymes slightly increased at 12h before dropping sharply, compared to OXPHOS-related enzymes that increased at 12h, stabilized until 24h and went back to their initial level. As observed by [37], at some point of the erythroid differentiation process, energy demand increased, and glycolysis as well as OXPHOS were enhanced. Moreover a peak of cell respiration was reported during erythroid differentiation at the high proliferation phase [46]. It has been shown that erythroid differentiation initiation indeed was characterized by an initial increase of the proliferation rate [16, 47]. Thus, the initiation of erythroid differentiation could be associated with an increase of the proliferation rates and of the energetic demand, involving both glycolytic and OXPHOS pathways. Such characteristic was also reported in nascent neural differentiation [48]. Similarly to our erythroid differentiation process, this study highlighted a metabolic exit event occurring within the first 24h of neural differentiation, corresponding to an increase of both glycolytic and mitochondrial parameters. Thus, both studies promote a new vision of metabolism behaviour during differentiation. The differentiation process is not only accompanied by a metabolic switch from one status straight to another, but it is rather characterized by metabolic pathways variations, mostly a peak of glycolytic flow, lactate generation and OXPHOS during the first 24h following differentiation induction [48]. Such occurrence might be seen as a mark of the exit from the stable self-renewing state and thus differentiation settlement.

It is tempting to relate this simultaneous increase in two somehow contradictory metabolic pathways with previous observations of: (1) a surge in gene expression variability at 8-24h of the differentiation process, preceding the irreversible commitment to differentiation [13]; and (2) the uncertainty phase at the cell level [49]. All these evidences of an unstable state, observable at the gene expression [13], metabolism [48] and phenotypic levels [49], preceding irreversible differentiation suggest that this variable behaviour might be a driving force for the differentiation process.

There are many know reasons why gene expression and metabolic variations might be involved in a correlative way in cell decision making in the context of differentiation. The most obvious link is through epigenetic modifications. Indeed, metabolites such as acetyl CoA generated from pyruvate and feeding OXPHOS, represent a substrate for histone acetylase [50]. On the other hand, the nicotinamide adenine dinucleotide (NAD), regenerated by various metabolic pathways, is involved in histone deacetylation. It was shown for instance that NAD concentration was proportional to the enzymatic kinetic of various histone deacetylases, including sirtuins and also other HDAC (histone deacetylases) [51]. Sirtuins are NAD-dependent histone-deacetylases that are intrinsically linked to the cellular energetic state through their dependence upon NAD [52]. Furthermore, various complexes involved in chromatin modification and remodelling are ATP consuming [53], which could participate in the peak of energy demand that we observed at the beginning of the differentiation process. Such a global remodelling could be an important aspect of the commitment process [13].

Therefore epigenetics could be the link between the concomitant gene expression and metabolism rearrangements [51, 54, 55] which might represent a circular causal driving force pushing the cells out of their self-renewing state and into their new differentiated state.

## Conclusion

We demonstrated that erythroid differentiation was accompanied with a metabolic switch from lactate-producing glycolysis toward OXPHOS, marked by an increased of ATP needs and both pathways; at 12h-24h of the differentiation process. Moreover our results suggested that LDHA could play an important role in erythroid progenitors self-renewal and its drop could be involved in the metabolic rearrangements occurring while they differentiate. Finally, we showed that LDHA decline and OXPHOS upkeep could be necessary for erythroid differentiation. Furthermore, our results are compatible with the vision of a differentiation process first passing through an hesitating phase allowing cells to escape from their stable self-renewing state, before committing to a differentiation pathway, toward a new stable state.

## Acknowledgments

We thank Andras Paldi for fruitful discussions about metabolism mechanisms complexity and critical reading. We also thank Ludivine Walter for her helpful insights. We acknowledge the contribution of the AniRA cytométrie en flux platform (Sébastien Dussurgey) of SFR BioSciences Gerland-Lyon Sud (UMS3444/US8) for cell sorting and training on the ISX device. We thank La Ligue National Contre le Cancer (Comité de Haute-Savoie) for funding, and the BioSyL Federation and the LabEx Ecofect (ANR-11-LABX-0048) of the University of Lyon for inspiring scientific events.

## References

1. Kim DY, Rhee I, Paik J. Metabolic circuits in neural stem cells. Cellular and Molecular Life Sciences. 2014;71(21):4221–4241. doi:10.1007/s00018-014-1686-0.

2. Israelsen WJ, Lee D, Yu VWC, Jeanson NT, Clish CB, Cantley LC, et al. Differential Dependence On Aerobic Glycolysis In Normal and Malignant Hematopoietic Stem and Progenitor Cells To Sustain Daughter Cell Production. Blood. 2013;122(21):793–793.

3. Shum LC, White NS, Mills BN, de Mesy Bentley KL, Eliseev RA. Energy Metabolism in Mesenchymal Stem Cells During Osteogenic Differentiation. Stem Cells and Development. 2015;25(2):114–122.

4. Zheng X, Boyer L, Jin M, Mertens J, Kim Y, Ma L, et al. Metabolic reprogramming during neuronal differentiation from aerobic glycolysis to neuronal oxidative phosphorylation. eLife. 2016;5:e13374. doi:10.7554/eLife.13374.

5. Sandoval IT, Delacruz RGC, Miller BN, Hill S, Olson KA, Gabriel AE, et al. A metabolic switch controls intestinal differentiation downstream of Adenomatous polyposis coli (APC). eLife. 2017;6:e22706. doi:10.7554/eLife.22706.

6. Folmes CDL, Terzic A. Metabolic determinants of embryonic development and stem cell fate. Reproduction, Fertility and Development. 2014;27(1):82–88. doi:https://doi.org/10.1071/RD14383.

7. Folmes CDL, Nelson TJ, Martinez-Fernandez A, Arrell DK, Lindor JZ, Dzeja PP, et al. Somatic Oxidative Bioenergetics Transitions into Pluripotency-Dependent Glycolysis to Facilitate Nuclear Reprogramming. Cell Metabolism. 2011;14, 264–271. doi:10.1016/j.cmet.2011.06.011.

8. Baron MH, Isern J, Fraser ST. The embryonic origins of erythropoiesis in mammals. Blood. 2012;119(21):4828–37. doi:10.1182/blood-2012-01-153486.

9. Xue QF, Yeung ES. Variability of Intracellular Lactate-Dehydrogenase Isoenzymes in Single Human Erythrocytes. Analytical Chemistry. 1994;66(7):1175–1178. doi:DOI 10.1021/ac00079a036.

10. Zail SS, Vandenhoek AK. Lactate-Dehydrogenase Isoenzymes of Human Erythrocyte-Membranes. Clinica Chimica Acta. 1977;79(1):15–19. doi:Doi 10.1016/0009-8981(77)90454-5.

11. Sandoval H, Thiagarajan P, Dasgupta SK, Schumacher A, Prchal JT, Chen M, et al. Essential role for Nix in autophagic maturation of erythroid cells. Nature. 2008;454(7201):232–U66. doi:10.1038/nature07006.

12. Stier A, Bize P, Schull Q, Zoll J, Singh F, Geny B, et al. Avian erythrocytes have functional mitochondria, opening novel perspectives for birds as animal models in the study of ageing. Frontiers in Zoology. 2013;10. doi:Unsp 33 10.1186/1742-9994-10-33.

13. Richard A, Boullu L, Herbach U, Bonnafoux A, Morin V, Vallin E, et al. Single-Cell-Based Analysis Highlights a Surge in Cell-to-Cell Molecular Variability Preceding Irreversible Commitment in a Differentiation Process. PLoS Biol. 2016;14(12):e1002585. doi:10.1371/journal.pbio.1002585.

14. Isern J, He Z, Fraser ST, Nowotschin S, Ferrer-Vaquer A, Moore R, et al. Single-lineage transcriptome analysis reveals key regulatory pathways in primitive erythroid progenitors in the mouse embryo. Blood. 2011;117(18):4924–34. doi:10.1182/blood-2010-10-313676.

15. Gautier EF, Ducamp S, Leduc M, Salnot V, Guillonneau F, Dussiot M, et al. Comprehensive Proteomic Analysis of Human Erythropoiesis. Cell Rep. 2016;16(5):1470–1484. doi:10.1016/j.celrep.2016.06.085.

16. Gandrillon O, Schmidt U, Beug H, Samarut J. TGF-β cooperates with TGF-α to induce the self-renewal of normal erythrocytic progenitors: evidence for an autocrine mechanism. The EMBO Journal. 1999;18(10):2764–2781. doi:10.1093/emboj/18.10.2764.

17. Gandrillon O, Samarut J. Role of the different RAR isoforms in controlling the erythrocytic differentiation sequence. Interference with the v-erbA and p135gag-myb-ets nuclear oncogenes. Oncogene. 1998;16(5):563–74.

18. R Core Team. R: A Language and Environment for Statistical Computing; 2013. Available from: http://www.R-project.org/.

19. Leduc M, Gautier EF, Guillemin A, Broussard C, Salnot V, Lacombe C, et al. Deep proteomic analysis of chicken erythropoiesis. bioRxiv. 2018. doi: 10.1101/289728.

20. Kim JH, Lee SR, Li LH, Park HJ, Park JH, Lee KY, et al. High cleavage efficiency of a 2A peptide derived from porcine teschovirus-1 in human cell lines, zebrafish and mice. PLoS One. 2011;6(4):e18556. doi:10.1371/journal.pone.0018556.

21. Ran FA, Hsu PD, Wright J, Agarwala V, Scott DA, Zhang F. Genome engineering using the CRISPR-Cas9 system. Nat Protoc. 2013 Nov;8(11):2281–308. doi:10.1038/nprot.2013.143. Epub 2013 Oct 24. 10.1038/nprot.2013.143

22. Geiser M, Cebe R, Drewello D, Schmitz R. Integration of PCR fragments at any specific site within cloning vectors without the use of restriction enzymes and DNA ligase. Biotechniques. 2001;31(1):88–+.

23. Wilson JE. Isozymes of mammalian hexokinase: structure, subcellular localization and metabolic function. Journal of Experimental Biology. 2003;206(12):2049–2057. doi:10.1242/jeb.00241.

24. Mor ECC I, Vousden KH. ontrol of glycolysis through regulation of PFK1: old friends and recent additions. Cold Spring Harb Symp Quant Biol. 2011;76:211–216.

25. Jurica MS, Mesecar A, Heath PJ, Shi WX, Nowak T, Stoddard BL. The allosteric regulation of pyruvate kinase by fructose-1,6-bisphosphate. Structure. 1998;6(2):195–210. doi:Doi 10.1016/S0969-2126(98)00021-5.

26. Patel MS, Roche TE. Molecular biology and biochemistry of pyruvate dehydrogenase complexes. FASEB J. 1990;4(14):3224–33.

27. Sugden MC, Holness MJ. Recent advances in mechanisms regulating glucose oxidation at the level of the pyruvate dehydrogenase complex by PDKs. American Journal of Physiology-Endocrinology and Metabolism. 2003;284(5):E855–E862. doi:10.1152/ajpendo.00526.2002.

28. Chan SK, Margoliash E. Amino acid sequence of chicken heart cytochrome c. J Biol Chem. 1966;241(2):507–15.

29. Nishikimi M, Hosokawa Y, Toda H, Suzuki H, Ozawa T. The Primary Structure of Human Rieske Iron-Sulfur Protein of Mitochondrial Cytochrome Bc1 Complex Deduced from Cdna Analysis. Biochemistry International. 1990;20(1):155–160.

30. Johnston IG, Gaal B, das Neves RP, Enver T, Iborra FJ, Jones NS. Mitochondrial Variability as a Source of Extrinsic Cellular Noise. Plos Computational Biology. 2012;8(3). doi:ARTN e1002416 10.1371/journal.pcbi.1002416.

31. Manerba M, Vettraino M, Fiume L, Di Stefano G, Sartini A, Giacomini E, et al. Galloflavin (CAS 568-80-9): a novel inhibitor of lactate dehydrogenase. ChemMedChem. 2012;7(2):311–7. doi:10.1002/cmdc.201100471.

32. Deck LM, Royer RE, Chamblee BB, Hernandez VM, Malone RR, Torres JE, et al. Selective Inhibitors of Human Lactate Dehydrogenases and Lactate Dehydrogenase from the Malarial Parasite Plasmodium falciparum. Journal of Medicinal Chemistry. 1998;41(20):3879–3887. doi:10.1021/jm980334n.

33. Le A, Cooper CR, Gouw AM, Dinavahi R, Maitra A, Deck LM, et al. Inhibition of lactate dehydrogenase A induces oxidative stress and inhibits tumor progression. Proc Natl Acad Sci U S A. 2010;107(5):2037–42. doi:10.1073/pnas.0914433107.

34. Galluzzi L, Kepp O, Vander Heiden MG, Kroemer G. Metabolic targets for cancer therapy (vol 12, pg 829, 2013). Nature Reviews Drug Discovery. 2013;12(12):965–965. doi:10.1038/nrd4191.

35. Zhang Y, Zhang X, Wang X, Gan L, Yu G, Chen Y, et al. Inhibition of LDH-A by lentivirus-mediated small interfering RNA suppresses intestinal-type gastric cancer tumorigenicity through the downregulation of Oct4. Cancer Letters. 2012;321(1):45–54. doi:https://doi.org/10.1016/j.canlet.2012.03.013.

36. Rellinger EJ, Craig BT, Alvarez AL, Dusek HL, Kim KW, Qiao J, et al. FX11 inhibits aerobic glycolysis and growth of neuroblastoma cells. Surgery. 2017;161(3):747–752. doi:https://doi.org/10.1016/j.surg.2016.09.009.

37. Oburoglu L, Tardito S, Fritz V, de Barros SC, Merida P, Craveiro M, et al. Glucose and glutamine metabolism regulate human hematopoietic stem cell lineage specification. Cell Stem Cell. 2014;15(2):169–84. doi:10.1016/j.stem.2014.06.002.

38. Mejia-Pous C, Damiola F, Gandrillon O. Cholesterol synthesis-related enzyme oxidosqualene cyclase is required to maintain self-renewal in primary erythroid progenitors. Cell Proliferation. 2011;44(5):441–452. doi:10.1111/j.1365-2184.2011.00771.x.

39. Jones W, Bianchi K. Aerobic glycolysis: beyond proliferation. Front Immunol. 2015;6:227. doi:10.3389/fimmu.2015.00227.

40. Wang YH, Israelsen WJ, Lee DJ, Yu VWC, Jeanson NT, Clish CB, et al. Cell-State-Specific Metabolic Dependency in Hematopoiesis and Leukemogenesis. Cell. 2014;158(6):1309–1323. doi:10.1016/j.cell.2014.07.048.

41. Shyh-Chang N, Ng HH. The metabolic programming of stem cells. Genes & Development. 2017;31(4):336–346. doi:10.1101/gad.293167.116.

42. Frauwirth KA, Thompson CB. Regulation of T lymphocyte metabolism. Journal of Immunology. 2004;172(8):4661–4665. doi:DOI 10.4049/jimmunol.172.8.4661.

43. Panopoulos AD, Yanes O, Ruiz S, Kida YS, Diep D, Tautenhahn R, et al. The metabolome of induced pluripotent stem cells reveals metabolic changes occurring in somatic cell reprogramming. Cell Res. 2012;22(1):168–77. doi:10.1038/cr.2011.177.

44. Rodriguez-Colman MJ, Schewe M, Meerlo M, Stigter E, Gerrits J, Pras-Raves M, et al. Interplay between metabolic identities in the intestinal crypt supports stem cell function. Nature. 2017;543(7645):424–+. doi:10.1038/nature21673.

45. Theret M, Gsaier L, Schaffer B, Juban G, Ben Larbi S, Weiss-Gayet M et al. AMPKα1-LDH pathway regulates muscle stem cell self-renewal by controlling metabolic homeostasis. The EMBO journal. 2017;36. doi:10.15252/embj.201695273.

46. Browne SM, Daud H, Murphy WG, Al-Rubeai M. Measuring dissolved oxygen to track erythroid differentiation of hematopoietic progenitor cells in culture. Journal of Biotechnology. 2014;187:135–138. doi:10.1016/j.jbiotec.2014.07.433.

47. Müllner E, Dolznig H, Beug H. Cell cycle regulation and erythroid differentiation. Curr Top Microbiol Immunol. 1996; p. 212:175–94.

48. Lees JG, Gardner DK, Harvey AJ. Mitochondrial and glycolytic remodeling during nascent neural differentiation of human pluripotent stem cells. Development. 2018;doi:10.1242/dev.168997.

49. Moussy A, Cosette J, Parmentier R, da Silva C, Corre G, Richard A, et al. Integrated time-lapse and single-cell transcription studies highlight the variable and dynamic nature of human hematopoietic cell fate commitment. Plos Biology. 2017;in the press.

50. Moussaieff A, Rouleau M, Kitsberg D, Cohen M, Levy G, Barasch D, et al. Glycolysis-mediated changes in acetyl-CoA and histone acetylation control the early differentiation of embryonic stem cells. Cell Metab. 2015;21(3):392–402. doi:10.1016/j.cmet.2015.02.002.

51. Reid MA, Dai Z, Locasale JW. The impact of cellular metabolism on chromatin dynamics and epigenetics. Nat Cell Biol. 2017;19(11):1298–1306. doi:10.1038/ncb3629.

52. Imai S, Johnson FB, Marciniak RA, McVey M, Park PU, Guarente L. Sir2: an NAD-dependent histone deacetylase that connects chromatin silencing, metabolism, and aging. Cold Spring Harb Symp Quant Biol. 2000;65:297–302.

53. Hota SK, Bruneau BG. ATP-dependent chromatin remodeling during mammalian development. Development. 2016;143(16):2882–97. doi:10.1242/dev.128892.

54. Yong CS, Moussa DA, Cretenet G, Kinet S, Dardalhon V, Taylor N. Metabolic orchestration of T lineage differentiation and function. FEBS Letters. 2017;591:3104—3118. doi:10.1002/1873-3468.12849.

55. Folmes CDL, Terzic A. Energy metabolism in the acquisition and maintenance of stemness. Seminars in Cell & Developmental Biology. 2016;52:68–75. doi:10.1016/j.semcdb.2016.02.010.

